# Recombinant T cell receptors specific for HLA-A*02:01-restricted neoepitopes containing KRAS codon 12 hotspot mutations

**DOI:** 10.1101/2020.06.15.149021

**Authors:** Craig M. Rive, Eric Yung, Christopher S. Hughes, Scott D. Brown, Govinda Sharma, Lisa Dreolini, Nasrin M. Mawji, Cassia Warren, Joanna M. Karasinska, Jonathan M. Loree, Donald T. Yapp, Gregg B. Morin, Daniel J. Renouf, David F. Schaeffer, Simon Turcotte, Robert A. Holt

## Abstract

KRAS codon 12 mutations are among the most common hotspot mutations in human cancer. Using a functional screening platform we set out to identify αβ T-cell receptors (TCRs) as potential targeting reagents for KRAS^G12D^ and/or KRAS^G12V^ neoepitopes presented by the prevalent HLA-A*02:01 allele. Here we describe isolation and characterization of three distinct CD8^+^ T cell clones from a pre-treated 76 year old patient with pancreatic ductal adenocarcinoma (PDAC). One clone was KRAS^G12V^ reactive and two clones were KRAS^G12D^ reactive. Tetramer staining showed high specificity of each T cell clone for its cognate HLA-A*02:01 restricted KRAS^G12V^ or KRAS^G12D^ neoepitope (>98% tetramer positive) without appreciable cross-reactivity to wild-type KRAS (<2% tetramer positive). We amplified and sequenced the full-length TCR alpha and beta chains from each of the three T cell clones and determined that these three TCRs comprised distinct combinations of two different TCR alpha chains and two distinct TCR beta chains. We resynthesized these TCR alpha and beta chain nucleotide sequences and reconstituted the original pairs in healthy donor CD8^+^ T cells by lentiviral transduction, substituting the human αβ TCR constant gene segments with murine αβ TCR constant gene segments to prevent mispairing with endogenous TCR subunits. Tetramer analysis and IFN-γ ELISpot analysis confirmed the specificity of each reconstituted TCR for its cognate HLA-A*02:01 restricted KRAS neoepitope. To test cytolytic activity TCR-transduced healthy donor CD8^+^ T cells were co-cultured with KRAS^G12V^, KRAS^G12D^ or KRAS^wt^ peptide-pulsed K562-HLA-A*02:01 antigen presenting cells at an effector to target cell ratio of 4:1. Under these conditions we observed neoepitope-specific killing of 16.5% to 19.0% of target cell populations. To assess *in vivo* activity we developed a KRAS^G12V^/A*02:01 patient-derived xenograft (PDX) mouse model. Over a 56-day period, PDX bearing mice infused with human TCR-transduced T cells had significantly reduced tumor growth and longer survival compared to mice infused with non-transduced control T cells. In conjunction with other therapeutic approaches, immune effector cell therapies expressing these TCRs may improve outcomes for HLA-A*02:01 patients with KRAS^G12V^ and/or KRAS^G12D^ positive tumors.

## Introduction

The p21/ras proteins are a family of highly conserved intracellular GTPases that mediate key cell proliferation and differentiation pathways via the RAS/MAPK pathway. Ras proteins typically acquire oncogenic potential as the result of single amino acid substitutions in codons 12, 13, or 61. KRAS codon 12 mutations are among the most common hotspot mutations in human cancer. They are particularly frequent in pancreatic ductal adenocarcinoma (PDAC), where they are also associated with poor prognosis (*1*). Over 90% of PDAC tumours are KRAS mutation positive with approximately 37% of tumours bearing a KRAS^G12D^ mutation and approximately 29% of tumours bearing a KRAS^G12V^ mutation. KRAS codon 12 mutations are also frequently observed in colorectal adenocarcinomas (13% KRAS^G12D^; 9% KRAS^G12V^) and adenocarcinomas of the lung (4% KRAS^G12D^; 6% KRAS^G12V^) where they are likewise associated with worse outcome (*2-7*). These mutation frequencies are from the GENIE consortium’s analysis of extensive clinical-grade tumour genome data (*8*), and recapitulate findings from earlier, large-scale tumour genome sequencing efforts (*9-11*) and independent studies (*12-15*). Codon 12 mutations in the KRAS protein cause it to become locked in an activated GTP-bound state, which constitutively activates growth signaling and increases the likelihood of oncogenic transformation of the mutated cells (*16*). Difficulty exploiting KRAS mutations pharmacologically has prompted exploration of this oncogene as an immunological target (*17-19*). It has been shown previously that KRAS codon 12 containing mutant peptides processed intracellularly and presented at the cell surface bound to MHC Class I receptors are capable of eliciting a cytolytic T cell response against the mutated cell (*20, 21*). *In silico* epitope prediction methods applied to comprehensive tumour mutation datasets support the possibility that highly recurrent KRAS hotspot mutations yield highly recurrent neoepitopes (*22-25*).

Recently, Wang *et al*. isolated restricted KRAS^G12D^ and KRAS^G12V^ reactive T-cell receptors (TCRs) from HLA-A*11:01^+^ transgenic mice and observed that human peripheral blood mononuclear cells (PBMCs) transduced with these TCRs could recognize HLA-A*11:01^+^ tumour lines bearing the canonical KRAS mutations. Adoptive transfer of PBMC tranduced with these TCRs significantly reduced tumour growth in an HLA-A*11:01, KRAS^G12D^ mutated pancreatic cell line xenograft model (*26*). Subsequently, Tran *et al*. isolated and expanded tumour infiltrating T cells reactive against an HLA-C*08:02-restricted KRAS^G12D^ neoepitope from a patient with metastatic colon cancer (*27*). Infusion of this autologous cell product as part of a clinical trial led to regression of the patient’s metastatic lung lesions, but loss of expression of the HLA-C*08:02 allele in one of the lesions allowed subsequent progression of disease (*27*). These findings are encouraging and support the notion that anti-tumour T-cell responses targeting HLA class I neoepitopes originating from mutant KRAS proteins are potentially therapeutically relevant.

We set out to identify TCRs that could be used to produce TCR-transduced T cell products for treating tumours possessing KRAS^G12D^ and/or KRAS^G12V^ neoepitopes presented by the highly prevalent HLA-A*02:01 allele. HLA frequency varies by population, with HLA-A*02:01 frequencies as high as 54.5% in North American populations, 34.4% in European populations, 40.5% in Asian populations, and 18.4% in African populations, according to study data compiled by the Allele Frequency Net Database (*28*). Here, we describe screening PBMC from a 76 year old HLA-A*02:01 patient diagnosed with a stage 2 primary PDAC tumour, arising at the pancreatic tail, with regional lymph node involvement but no distant metastasis. PBMC used for TCR discovery were obtained post-operatively, prior to chemotherapy and radiation therapy that were delivered in the adjuvant setting. From this subject we isolated three HLA-A*02:01 restricted T cell clones; two specific for the KRAS^G12D^ amino acid 5-14 epitope (KLVVVGADGV) and one specific for the KRAS^G12V^ amino acid 5-14 epitope (KLVVVGAVGV). We amplified, sequenced, and re-synthesized the alpha and beta TCR chains from these T cell clones and reconstituted them in HLA-A*02:01 healthy donor CD8^+^ T cells. These TCR-modified healthy donor T cells recapitulated the epitope specific binding and cytolytic activity of the original clones and showed efficacy in a PDAC patient-derived xenograft (PDX) tumour model.

## Results

We first confirmed that the KRAS 5-14 peptide could be naturally processed and presented. We used PANC-1 cells which are heterozygous for both the KRAS^G12D^ mutation and the HLA-A*02:01 allele (*29*). The mutant peptide was detectable by surface elution and targeted Multiple Reaction Monitoring (MRM) mass spectrometry (**Figure S1**), although at a lower level than the KRAS^wt^ peptide.

Next, to isolate neoantigen-specific CD8^+^ T cells we deployed a “miniline” functional screening platform that we previously established to evaluate the T cell response to High Grade Serous Ovarian Cancer (*30,31*). Briefly, polyclonal CD8^+^ T cells were magnetically sorted from PBMC, distributed in 96 well plates at a density of several thousand T cells per well, expanded non-specifically, rested, and then screened by interferon*-*gamma (IFN-γ) ELISpot for reactivity against peptide-pulsed K562-HLA*02:01 artificial Antigen Presenting Cells (aAPC). T cells from positive pools that showed up-regulation of the CD137 activation marker when co-incubated with peptide-pulsed aAPC were sorted into individual wells of a 96 well plate and clonally expanded. Using this approach we isolated one HLA-A*02:01-restricted KRAS^G12V^ specific T cell clone and two HLA-A*02:01-restricted KRAS^G12D^ T cell clones. We were somewhat surprised to recover T cell clones reactive to both KRAS^G12V^ and KRAS^G12D^ neoepitopes from a single subject. These mutations have previously been reported to co-occur in some patients (*32*), but we did not have access to tumour material to determine the mutational status of this particular subject. However, HLA class I typing of the three T cell clones verified they are matched (HLA-A*02:01, -A*01:01, -B*07:02, -B*44:02, - C*07:02, and -C*05:01) providing assurance these T cells do in fact originate from the same patient.Tetramer staining showed high specificity of each T cell clone for its cognate HLA-A*02:01-restricted KRAS neoepitope (**Figure 1**). Specifically, 99.0 +/- 0.1% (mean +/- SD) of the HLA-A*02:01-restricted, KRAS^G12V^-specific monoclonal T cells stained positive for the KRAS^G12V^ - A*02:01 tetramer. The two HLA-A*02:01-restricted, KRAS^G12D-^specific monoclonal T cells stained 99.3 +/- 0.1% and 98.8 +/- 0.9% positive for the KRAS^G12D^ - A*02:01 tetramer, respectively. Each T cell clone stained <2% positive for KRAS wild-type HLA-A*02:01 tetramer. Of note, we initially attempted to isolate KRAS neoeptiope reactive CD8^+^ T cells directly from patient PBMC by tetramer sorting, but these attempts did not lead to successful expansion of any T cell clones, necessitating our adoption of the more comprehensive miniline-based search strategy described here.

**Figure 1:**
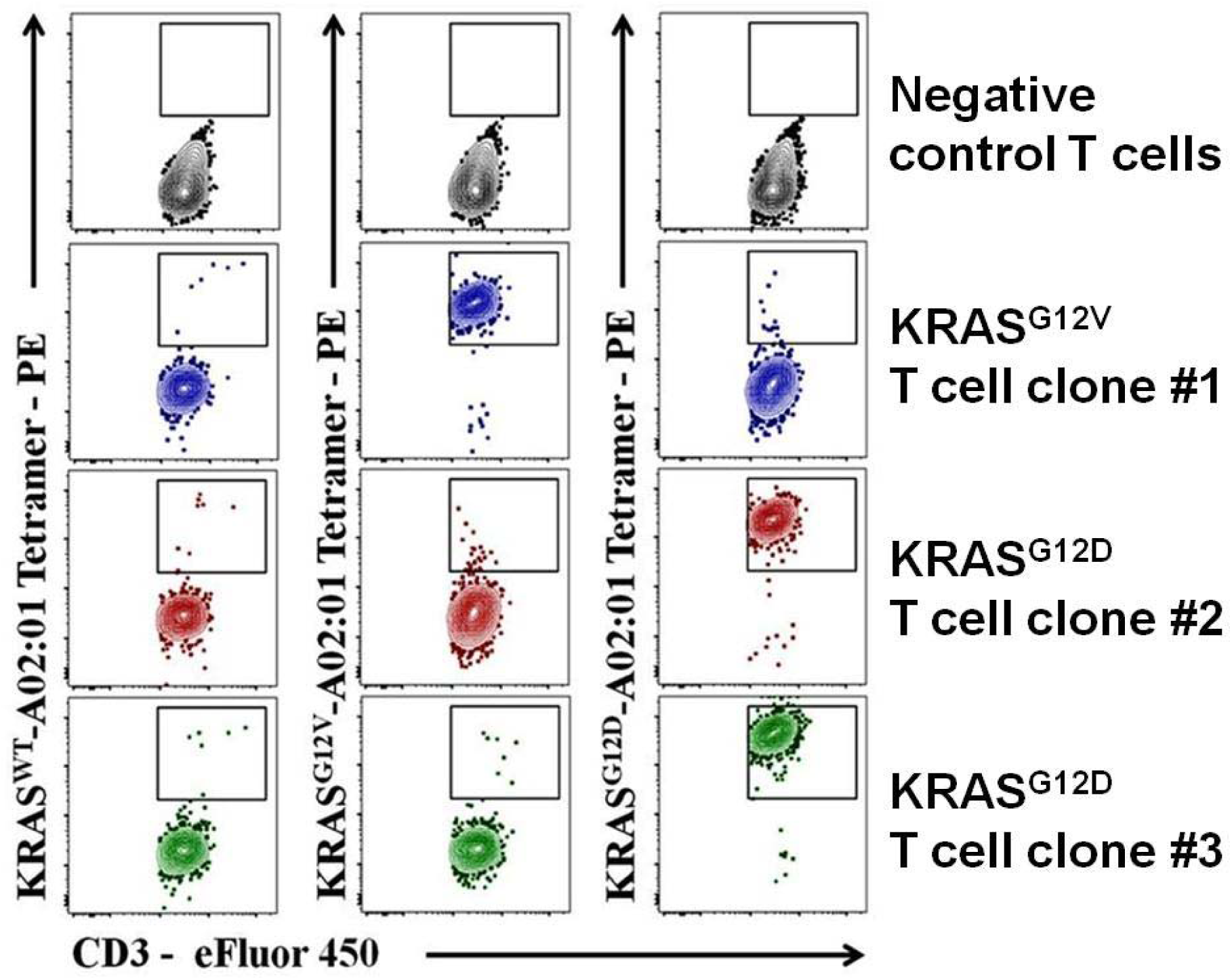
Tetramer analysis of HLA-A*02:01-restricted KRAS^G12V/D^ specific T cell clones. Monoclonal CD8+ T cells (blue) were stained strongly by the KRAS^G12V^/A*02:01 tetramer (99.1%) and minimally (<2%) by the KRAS^G12D^ tetramer and KRAS^wt^ tetramer. The two other monoclonal CD8+ T cells (red and green) stained strongly by the KRAS^G12D^/A*02:01 tetramer (98.8% and 97.5%, respectively) and minimally (<2%) by the KRAS^G12V^ and KRAS^wt^ tetramers. The flow gating protocol is outlined in Figure S3.

Next, we amplified and sequenced (*33*) the full-length TCR alpha and beta chains from each of the three T cell clones which revealed two distinct alpha chains and two distinct beta chains shared among the three clones **(Table 1)**. The nucleotide sequences encoding the cognate TCR alpha and beta chain pair for each T cell clone were synthesized and inserted, in the configuration Alpha-T2A-Beta-P2A-mStrawberry, into a lentiviral transfer plasmid (*34*), under control of the human EF1α promoter. *The synthetic TCRs corresponding to each of T cell clones 1, 2 and 3 are referred to hereafter as KTCR1, KTCR2, and KTCR3, respectively*. In each of these synthetic TCRs the human constant gene segment of each chain was replaced with the orthologous murine constant gene segment to facilitate correct alpha-beta chain pairing (*36*). Replication-incompetent lentivirus was generated for each KTCR construct and used to transduce CD8^+^ T cells obtained from HLA-A*02:01^+^ normal healthy donors, followed by sorting and expansion of successfully transduced T cells from each population.

**Table 1:** Neoepitope specificities and alpha/beta chain identities of TCRs 1, 2 and 3 (derived from T cell clones 1, 2 and 3, respectively).

| **TCR $\alpha\beta$ variable chains** | **TRBV 19\*01** | **TRBV 04-1\*01** |
| --- | --- | --- |
| **TRAV 27\*01** | TCR1 (KRAS<sup>G12V</sup>) | TCR3 (KRAS<sup>G12D</sup>) |
| **TRAV 13-2\*01** | TCR2 (KRAS<sup>G12D</sup>) |  |

Tetramer analysis of TCR-transduced T cells mirrored the results observed from tetramer analysis of the original T cell clones. Specifically, 83.7 +/- 0.5% (mean +/- SD) of KTCR1-transduced CD8^+^ T cells stained positive for the KRAS^G12V^ - A*02:01 tetramer, 89.3 +/- 0.5% of KTCR2-transduced CD8^+^ T cells stained positive for the KRAS^G12D^ - A*02:01 tetramer and 88.7 +/- 0.5% of KTCR3-transduced CD8^+^ T cells likewise stained positive for the KRAS^G12D^ - A*02:01 tetramer. Each population of KTCR transduced T cells stained < 5% positive for the KRAS wild-type HLA-A*02:01 tetramer (**Figure 2**).

**Figure 2:**
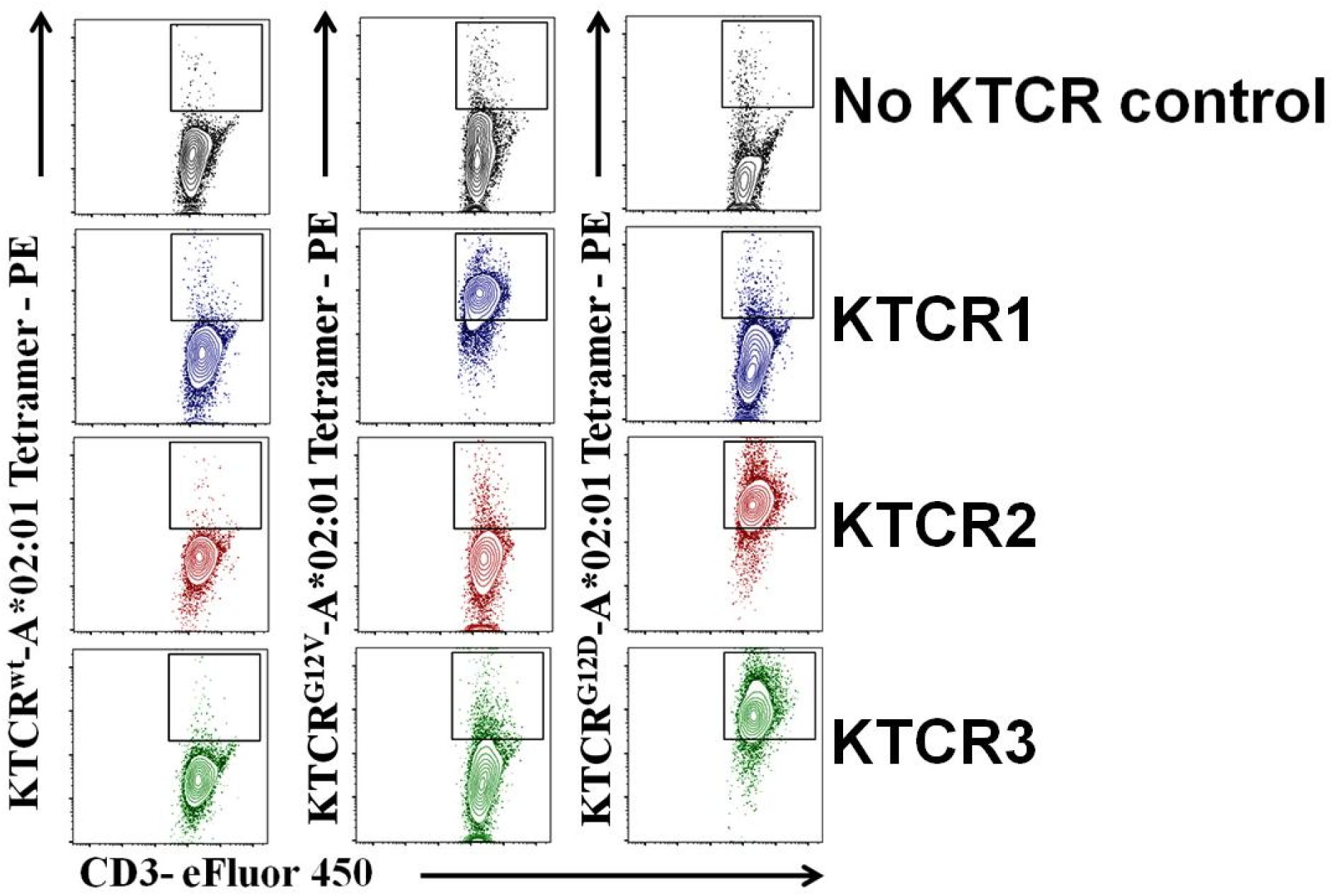
Tetramer analysis of KTCR-transduced CD8+ T cells. KTCR1 (blue) transduced CD3+CD8+ T cells (blue) were stained strongly by the KRAS^G12V^/A*02:01 tetramer (83.7%) and minimally (<5%) by KRAS^G12D^ and KRAS^wt^ tetramers. KTCR2 (red) and KTCR3 (green) transduced CD3+CD8+ T cells were stained strongly by the KRAS^G12D^/A*02:01 tetramer (89.3% and 88.7%, respectively) and minimally (<5%) by KRAS^G12V^ and KRAS^wt^ tetramers. The flow gating protocol is outlined in Figure S3.

Further verification of specificity was obtained by IFN-γ ELISpot analysis (**Figure 3A**) using HLA-A*02:01 transduced K562 aAPCs pulsed with the KRAS^G12D/V^ peptides. KTCR1-transduced CD8^+^ T cells produced 734 +/- 157 IFNγ spot forming units (SFU) per million cells when co-cultured with KRAS^G12V-^peptide-pulsed K562-A*02:01 target cells (50,000 cells each at a ratio of 1:1), compared to KRAS^G12D-^ and KRAS^wt-^pulsed target cells, which produced 31 +/- 47 and 26 +/- 43 SFU/million cells, respectively (p < 0.001). The KTCR2-transduced CD8^+^ T cells produced 360 +/- 77 IFNγ SFU per million cells when co-cultured with KRAS^G12D^-peptide-pulsed target cells, compared to KRAS^G12V-^and KRAS^wt^-pulsed target cells which produced 26 +/- 40 and 26 +/- 52 SFU/million cells,\ respectively (p < 0.001). KTCR3-transduced CD8^+^ T cells produced 647 +/- 162 IFNγ SFU/million cells when co-cultured with KRAS^G12D^-peptide-pulsed target cells compared to KRAS^G12V-^ and KRAS^wt^-pulsed target cells which produced 87 +/- 62 and 29 +/- 28 SFU/million cells, respectively (p < 0.001 and p < 0.002, respectively) (**Figure 3A**).

**Figure 3:**
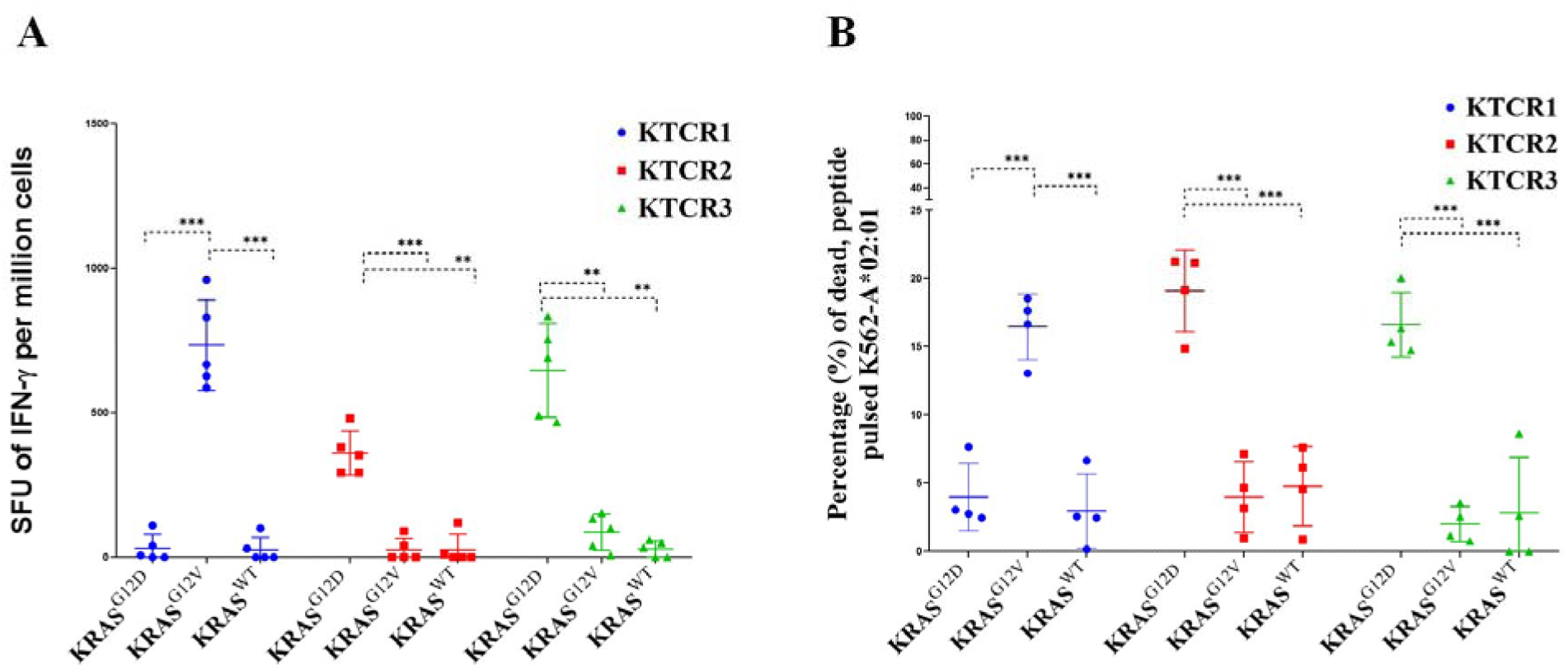
*In vitro* analysis of KTCR transduced CD8^+^ T cells. **Results from** IFN-γ ELISpot assays (A) and cytolytic assays (B), evaluating KTCR1 (blue circles), KTCR2 (red squares), and KTCR3 (green triangles) activity against peptide pulsed K562-A*02:01 target cells. (**p < 0.01 and ***p = 0.001; ANOVA and Tukey’s multiple comparison test)The flow gating protocol is outlined in **Figure S4**.

The function of KTCR1-, KTCR2-, or KTCR3-transduced CD8^+^ T cells was evaluated using cytotoxic assays. Transduced T cells were co-cultured for 5 hours with peptide-pulsed K562-HLA-A*02:01 target cells at an effector to target cell ratio of 4:1. Killing of aAPC was measured using vital staining and flow cytometric analysis. The KTCR1-transduced CD8^+^ T cells killed 16.5 +/- 2.41% (mean +/- SD) of the KRAS^G12D-^peptide-pulsed target cells compared to 4.0 +/- 2.5% of KRAS^G1V-^ and 2.9 +/- 2.7% of KRAS^WT-^ pulsed target cells. The KTCR2-transduced CD8^+^ T cells killed 19.1 +/- 3.0% of KRAS^G12D-^ pulsed target cells compared to 4.0 +/- 2.6% of KRAS^G12V-^ and 4.8 +/- 2.9% of KRAS^WT-^pulsed target cells (p <0.001). KTCR3-transduced CD8^+^ T cells killed 16.6 +/- 2.4% of KRAS^G12D-^pulsed target cells compared to 4.0 +/- 2.6% of KRAS^G12V-^ and 4.8 +/- 2.9% of KRAS^WT-^pulsed target cells (p <0.001) (**Figure 3B**).

KTCR1 was assessed *in vivo* using a PDX mouse model. Initially, we screened a number of PDX samples in order to find one which possessed both the KRAS^G12V^ mutation and the HLA-A*02:01 allele. Once a suitable PDX sample was identified, tumours were expanded by passaging them in CB17SC-M mice. Expanded PDX tumours were then surgically implanted into experimental CB17SC-M mice and once tumours were established and had reached an average size of 100mm^3^ the mice were infused with either KTCR1-transduced CD8^+^ T cells or with an equivalent dose of control CD8^+^ T cells expanded from the same donor but lacking KTCR1 expression. The reasoning behind treating mice with established tumours rather than early stage tumours was to better emulate a clinical setting. Infusion of the KTCR1-transduced CD8^+^ T cells on day 25 led to reduced tumour growth almost immediately post infusion, relative to control mice, with a statistically significant reduction at day 44 (p ≤ 0.02) and beyond (p ≤ 0.002 at Day 56) (**Figure 4A**). We humanely euthanized mice once the tumours reached a size of 1000mm^3^ and used this as a proxy for survival. The mice treated with the KTCR1-transduced CD8^+^ T cells showed longer survival compared to mice treated with CD8^+^ T cells that did not express KTCR1 (p ≤ 0.05) (**Figure 4B**).

**Figure 4:**
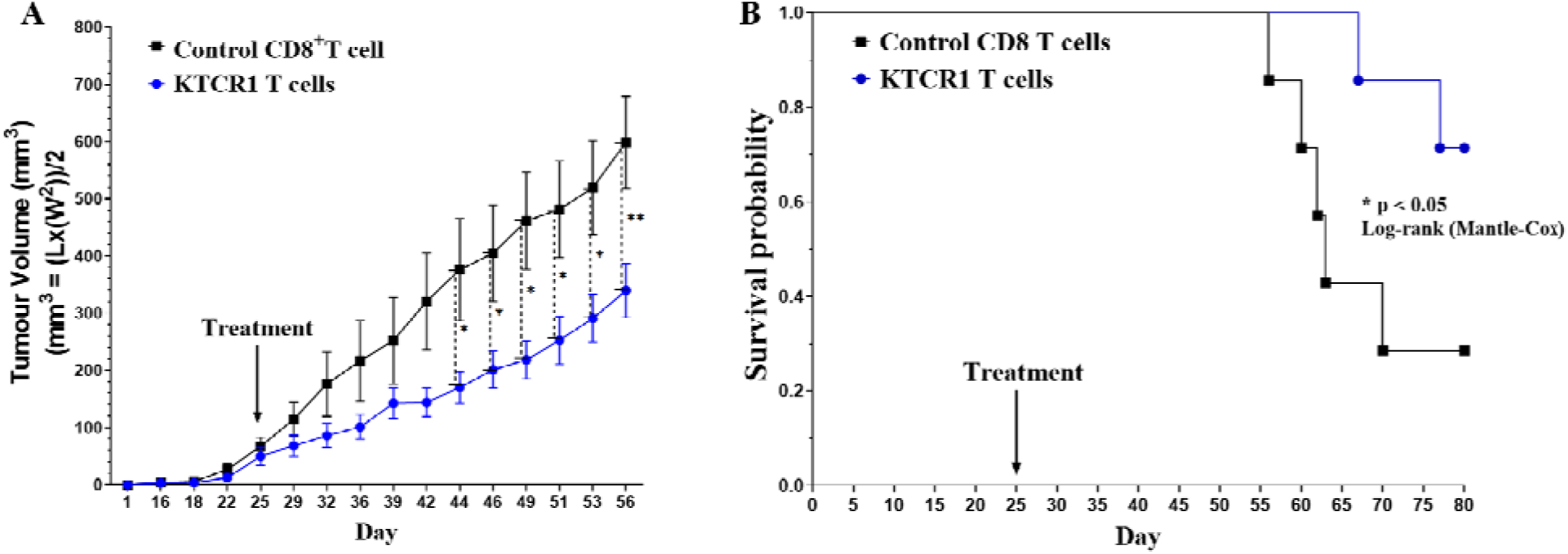
Testing of HLA-A*02:01-restricted KRAS^G12V^ specific TCR reconstituted T cells *in vivo*. **Figure 4A**: Treatment with KTCR1-transduced CD8^+^ T cells (blue circles) reduced tumour growth when compared to the mice treated with control CD8^+^ T cells (black squares) (n=7). Significantly reduced growth was seen from days 44 (* p ≤ 0.020), to completion of the experiment on day 56 (**p ≤ 0.002) (ANOVA, p < 0.001 and Tukey’s multiple comparison test). **Figure 4B**. Kaplan-Meier analysis of survival probability of mice treated with KTCR1 CD8^+^ T cells (blue circles) compared to mice treated with the control CD8^+^ T cells (black squares) (n=7; *p<0.05; mantel-Cox log-rank test).

## Discussion

We have isolated, from a newly diagnosed 76 year old PDAC patient, three distinct αβTCRs specific for HLA-A*02:01-binding KRAS^G12V^ and KRAS^G12D^ neoepitopes.

Interestingly, we observed that the HLA-A*02:01-restricted KRAS^G12V^ specific KTCR1 and HLA-A*02:01-restricted KRAS^G12D^ KTCR2 shared the same beta chain, suggesting that mutation specific binding was conferred by the alpha chain. However, we also observed that HLA-A*02:01-restricted KRAS^G12V^ specific KTCR1 and HLA-A*02:01-restricted KRAS^G12D^ specific KTCR3 shared the same alpha chain, which would suggest the opposite, that the beta chain confers mutation specificity **(Table 1)**. The simplest explanation for these observations would be that these were not pure clones but rather mixed populations. However, upon careful inspection, the sequence data indicated no polyclonality and HLA class I typing of the individual T cell clones confirmed that they did in fact come from the same patient. The existance of natural, mixed-chain TCRs with distinct specificities suggests that engineering therapeutic TCRs with novel subunit combinations could be beneficial.

Historically, there have been many efforts to exploit recurrent KRAS somatic point mutations as pharmacological targets, but mutant KRAS remains largely “undruggable”. Much recent attention has been given to the development of KRAS^G12C^ inhibitors, a number of which are currently in various stages of development and testing. One of these, AMG150 (37-41), was shown to produce durable regression of tumours in mice, even if KRAS^G12C^ is expressed heterogeneously. It was also shown in a murine model that inhibiting the actions of the activated KRAS^G12C^ mutant, in the presence of the PD-1 inhibitor pembrolizumab, led to the clearance of KRAS^G12C^ tumours and subsequent rejection of KRAS^G12D^ tumour grafts, substantiating these codon 12 mutations as neoantigenic.

There is potential to exploit KRAS^G12V-^ and KRAS^G12D^-specific recombinant TCRs in engineered Immune Effector Cell (IEC) therapies. Similar to the Chimeric Antigen Receptor (CAR) T-cell therapeutic approach, a patient’s autologous T-cells could be genetically modified to express the relevant KTCR, followed by *ex vivo* cell expansion and re-infusion. This method can generate a large number of antigen specific T cells with the potential to eliminate tumour cells and impart immune memory against neoantigens originating from this potent oncogene. In this manner, an IEC therapy deploying these TCRs, alone or in conjunction with standard therapies, could improve outcomes for cancer patients that carry the common HLA-A*02:01 allele and the common KRAS^G12V^ or KRAS^G12V^ mutations.

## Materials and Methods

### Sample acquisition and storage

Patient samples were obtained from the BC Gastrointestinal and Pancreatic (GIP) Biobank, a repository of tissue and blood samples donated by patients with pancreatic disease. Control T cells were isolated from from leukapheresis products from healthy donors (STEMCELL Technologies, Vancouver, BC, Canada). This study was approved by the Research Ethics Board of the BC Cancer Agency/University of British Columbia (certificates H16-00291-A021, H19-01738). PBMCs from the patient samples and healthy donor were isolated by Ficoll gradient purification as per manufactures protocol (Ficoll, GE-17-1440, GE Healthcare). The isolated PBMC were stored by resuspending cells in cryopreservation media in aliquots of 5-10×10^6^ cells per ml and storaed on liquid nitrogen.

### Patient details

The patient was a 76 year old woman who was diagnosed after initial imaging identification and curative intent resection. The pathological diagnosis was that the patient had an invasive, well to moderately differentiated pancreatic adenocarcinoma, arising from an intraductal papillary mucinous neoplasm (IPMN), that exhibited regional lymph node metastases. The PBMC sample from which the TCRs were identified was acquired at the time of diagnosis and prior to any treatment. Post surgery, the patient received 3 cycles of adjuvant gemcitabine, then concurrent chemoradiotherapy with capecitabine and radiation therapy (50 Gy in 25 fraction) due to a positive margin. The patient then completed another 5 cycles of gemcitabine. Approximately 23 months after the original diagnosis, the patient experienced disease recurrence and passed away 14 months after the recurrence diagnosis.

### Cell culture media

All cell cultures were maintained in either a RPMI-1640 (Invitrogen, 11875-119) supplemented media or a DMEM (Invitrogen, 11995-073) supplemented media. RPMI-1640 supplemented media consisted of 2 mM GlutaMAX (Invitrogen, 35050-061), 1 mM MEM non-essential amino acid (Invitrogen, 11140-050), 1 mM sodium pyruvate (Invitrogen, 11360-070), 10 mM HEPES (Invitrogen, 15630-808), 100 U/mL penicillin/streptomycin (Invitrogen, 15140-122), MycoZap prophylactic (Lonza, VZA-2031), and 10% heat-inactivated fetal bovine serum (HI-FBS, Invitrogen, 12484028). DMEM supplemented media consisted of 2 mM GlutaMAX, 1 mM MEM non-essential amino acid, 100 U/mL penicillin-streptomycin, MycoZap prophylactic, and 10% HI-FBS. We used an in house MACS separation buffer comprised of D-PBS (Gibco, 59331C) with 2mM EDTA (ThermoFisher, 15575020) and 2% HI-FBS. FACS media consisted of D-PBS with 2% HI-FBS. Cryopreservation media consisted of HI-FBS with 10% DMSO, (Fisher Scientific, BP231-100). All media and supplements were filtered through a 0.22µM filter (VWR, 28199-730) and stored at 4°C. All cultures were maintained at 37°C and 5% CO_2_ atmosphere.

### Isolation of T cell clones

Screening patient T cells entailed a modified “mini-line” culture method, previously described by Martin *et al*. and Wick *et al*. (*30, 31*). CD8^+^ T cells were isolated from the patient PBMC sample using the Miltenyi MACS human CD8^+^ isolation kit, following the manufacturer’s protocol (Miltenyi, 130-096-495) and using an in house MACS separation buffer. The isolated CD8^+^ T cells were then centrifuged at 400xg for 10 min (Eppendorf Centrifuge 5810R, rotor A-4-62) at room temperature, the supernatant was removed, and the cell pellet resuspended in sRPMI-1640 media. Cell numbers were obtained using the Countess automated cell counter (Thermofisher). The cells were then centrifuged and the cell pellet resuspended in RPMI-1640 supplemented media with 300U/mL rhIL-2 (Peprotech, 200-02 and StemCell, 78036.3), 100 ng/mL of anti-CD3 (eBiosciences, 16-0037-85), 100 ng/mL anti-CD28 (BioLegend, 302943) soluble antibodies at a concentration of 20,000 cells/ ml. 100μl (2000 cells) were then plated into individual wells within a 96-well U-shaped plate (Thermo Fisher Scientific, 08-772-54) with an excess of allogeneic irradiated PBMCs feeders (2×10^5^ cells irradiated with 50 Gy, per 2000 T cells). Cells were cultured for 14 days and then were split into replicate plates on days 5, 7, 9, 11, and 13 of the culture. Fresh RPMI-1640 supplemented media with rhIL-2 was added to the split cultures, giving a final volume of 200 μl and a final rhIL-2 concentration of 300 IU/ml. On day 14, cells were washed with RPMI-1640 supplemented media and then rested for 2-4 days in RPMI-1640 supplemented media with 10 IU/ml rhIL-2 before each individual well was screened for KRAS^G12D/V^ peptide reactivity. Reactive T cells were single-cell sorted by Fluorescence Activated Cell Sorting (FACS) (Figure S2) based on detection of 4-1BB (CD137) expression, after being re-stimulated for 24 hours with HLA-A*02:01 positive aAPCs pulsed with up to 10µg/mL of the relevant KRAS^G12D/V^ peptides. After the co-culture, cells were stained with the surface staining antibodies, CD8-FITC and 4-1BB-CD137 APC (1/100, BioLegend). Cells were incubated at 37°C for 10 min, then washed by adding 2mL of FACS media to each tube and centrifuging for 10 minutes at 400xg at room temperature. The supernatant was discarded before resuspension in 500μl of FACS media with PI (1/1000, Invitrogen, P1304MP). Acquisition was performed on the BD LSRFortessa cell analyser, and analyzed using FlowJo and GraphPad Prism and sorted using BD FACS Aria. The monoclonal T cells were sorted into single wells of a 96-well U-shaped plate and expanded in RPMI-1640 supplemented media with rhIL-2 (300U/mL) and an excess of allogeneic irradiated PBMCs feeders (2×10^4^ cells irradiated with 50 Gy, per single T cells). The expanded candidate T cell clones were either resuspended in cryopreservation media and stored in liquid nitrogen or resuspended in RNAlater (invitrogen, AM7020) and stored as per the manufactures recommendations.

### HLA-class I typing

Candidate T cell clones were subjected to class I HLA sequencing and typing to verify identity. DNA was extracted from the T cells using the DNeasy Blood and Tissue kit (Qiagen, 69504) following the manufacturer’s protocol. To amplify the HLA-A, HLA-B, and HLA-C genes, PCR was performed using primers for HLA-A (forward 5-GAAACSGCCTCTGYGGGGAGAAGCAA and Reverse 5-TGTTGGTCCCAATTGTCTCCCCTC), HLA-B (forward 5-GGGAGGAGCGAGGGGACCS CAG and Reverse 5-GGAGGCCATCCCCGGCGACCTAT), and HLA-C (forward 5-AGCGAGG KGCCCGCCCGGCGA and Reverse 5-GGAG ATGGGGAAGGCTCCCCACT). The PCR assays were run using LA Taq polymerase (Takara Bio, TAKRR002) with the following thermocycling conditions: one cycle at 94°C for 4 min; 28 cycles at 98°C for 10 sec and 72°C for 4 minutes before cooling down and being held at 10°C. DNA from each sample was purified using a 1% agarose SYBR Safe e□Gel (Invitrogen, G442001) and a unique band between 850-1000bp size was excised and extracted using QIAGEN Multiple gel extraction kit (QIAGEN, 28704) before being resuspended in RNase/DNase□free water (Invitrogen, AM9916). The HLA-A, HLA-B, and HLA-C amplicons from the three T cell clones plus a control PBMC source were “A-tailed” by adding adenosine to the 3’ end of the amplicons. The amplicons where then ligated into the pCR4-topo plasmid between the M13F and M13R sequences and transformed into DH5α competent cells following the manufacturer’s protocols (Invitrogen, 18265-017). Up to 8 colonies were selected for HLA-A, HLA-B, and HLA-C sequencing for each T cell clone and the control sample and DNA was isolated using a QIAprep Spin Miniprep Kit (QIAGEN, 27104). Sanger sequencing was performed using the M13F and M13R primers (Genewiz). Sequences were then matched and blasted using IMGT and NCBI-IgBLAST databases to identify the HLA-typing of each T cell clone (*42-44*).

### Mass spectrometry

For MRM assay design, the HLA-A*02:01-restricted KRAS^wt^ (KLVVVGAGGV) and KRAS^G12D^ (KLVVVGADGV) aa5-14 peptides were synthesized (Peptide 2.0) and analyzed on an Orbitrap Fusion (Thermo Scientific) mass spectrometer (MS) using a data-dependent method with MS/MS scanning in the ion trap detector. Peptides were injected and chromatographically separated using an Easy nLC 1000 system (Thermo Scientific) with a trapping-analytical column packed in-house in 75µm inner-diameter (ID) fused silica capillaries using 3µm Reprosil-Pur C18 beads to lengths of 3 and 25cm, respectively. The analytical column was heated to 45°C using nanoSLEEVE ovens (Analytical Sales and Services). Survey scans (MS1) were acquired with a resolution of 120,000, a 350 – 1200 m/z mass range, 32ms maximum fill time, and a 200,000 automatic gain control (AGC) target. MS2 acquisition used a 1.6m/z isolation window, a fixed first mass of 110m/z, a higher-energy collisional dissociation energy of 30, rapid scan mode, a 60ms max fill time, and a 10,000 AGC target. Data was acquired in 60-minute runs using a 3-second maximum cycle for the above scan modes. Acquired MS2 scans were identified using Proteome Discoverer (version 1.4) by searching against the human proteome database (Uniprot, version 2014_10) (45) with the settings: precursor mass error 20ppm, fragment mass error 0.5 Daltons, carbamidomethylation as a fixed modification, and oxidation of methionine as a variable modification. Results files were imported into Skyline software and a total of 5 precursor-product ion MRM transitions were selected for specific detection of each of the wild-type and mutant peptides based on ranked intensities of the product ions.

Prior to analysis, MHC molecules were disassociated from the surface of 2×10^8^ PANC-1 cells by weak acid elution (0.131M citric acid, 0.066M Na2HP04, pH3.0). Cells were centrifuged at 400xg for 5 mins, (Eppendorf Centrifuge 5810R, rotor A-4-62) at room temperature. The supernatant was collected and passed over Amicon ultrafiltration device (Millipore Sigma, UFC9010, 10kDa cutoff) at 3200xg for 15 min. The resulting pool of epitopes were chemically labeled using reductive demethylation following previously established protocols (*47*). Synthetic peptides were processed using the same protocol, substituting deuterated formaldehyde to generate a ‘medium’ sample label. Labeling reactions were quenched by adding 1M glycine and incubating for 15-minutes at room temperature. Labeled samples were acidified using triflouroacetic acid (TFA) and cleaned-up using StageTip (3-disc plug of C18 Empore material, 3M). StageTip’s were conditioned by rinsing them with (2x) 0.1% TFA in acetonitrile and (2x) 0.1% TFA in water. Final elution was performed with two steps of 0.1% TFA in 60% acetonitrile. All steps were performed with 100uL volumes and at a flow rate of 1 drop-per-second using a vacuum manifold unit (Waters). Labeled and desalted peptides were concentrated in a SpeedVac and dried samples reconstituted in 1% TFA prior to MS analysis. Light labeled MHC eluted samples were spiked with medium synthetic peptides based on optimised concentrations, determined throughout the MS analysis.

Expressed epitopes presented by PANC-1 cells (ATCC® CRL-1469^™^) were analyzed on a QTRAP 6500 MS system coupled to an Eksigent nanoLC 415 system (AB Sciex) controlled by Analyst software (version 1.6.2). MS data acquisition was performed in non-scheduled MRM mode using a ‘High’ CAD gas setting, transition-specific collision energies, and a dwell time of 45ms per transition. A total of 5 precursor-product ion MRM transitions were used for specific detection of each of the wild-type and mutant peptides (**Table S1**). All MRM MS analyses were carried out using 30-minute runs at a chromatography flow rate of 0.45µL per minute. MHC elutions were performed in triplicate and samples analyzed individually by MRM-MS. Resultant data were processed in Skyline software (*46*) to yield area values for each peptide that were used for calculations of peptide amounts based on established concentrations of the spiked synthetic peptide.

### Interferon-gamma ELISpot assays

The panel of polyclonal T cell pools was then screened for reactivity to KRAS^G12D^, KRAS^G12V^ and KRAS^w/t^ peptides using IFN-γ ELISpot assays following the manufacturer’s protocol (MabTech). Briefly, ELISpot plates were coated with IFN-γ capture antibody (2 μg/ml per well, Mabtech mAb 1-D1K, 3420-7), in D-PBS and then blocked with RPMI-1640 supplemented media. At an effector to target ratio of 1:1, 1×10^5^ effector CD8^+^ T cells were incubated with 1×10^5^ aAPCs transduced to express HLA-A*02:01, pulsed with the appropriate KRAS^G12V/D^ peptide (0.1-100 µg/mL). Controls included 1×10^5^ effector CD8^+^ T cells co-cultured with anti-human CD3 monoclonal antibody (0.5 µg/mL, Mabtech, 3605-1-1000), or with media alone. ELISpot plates were then incubated at 37°C with 5% CO_2_ for 18 to 22 hours, then wells were washed 5 times with sterile D-PBS and incubated with anti–IFNγ biotinylated antibody (1 µg/mL per well, Mabtech, mAb 7-B6-1, 3420-9H) for 2 hours.

The ELISpot plates were washed as described above, and incubated for 1 hour with streptavidin-HRP (diluted 1:100 with D-PBS, Mabtech, 3310-9-1000), and then washed again. Plates were substrated for 8-12 minutes using 0.5µM filtered 3’3’5’5’-Tetramethylbenzidine (TMB substrate) (Mabtech, 3652-F10) and the plates were evaluated with an AID automated microplate ELISpot reader and associated software (AID ELISpot Version 7.0).

### Flow cytometry

HLA-A*02:01 - KRAS^G12V/D/wt^ tetramer staining assays involved staining both monoclonal T cells and KTCR expressing cells (**Figure 2**). Tetramers were manufactured by the NIH Tetramer core facility and labelled with the PE fluorochrome. Cells were stained with the tetramer plus CD3-eFluor 450 (1/100, 48-0032-82 eBioscience) and CD8-FITC (1/100, 300905 BioLegend) at 4°C for 30 min. Samples were then washed by adding 2mL of FACS media to each tube and centrifuging for 10 minutes at 400xg at 4°C. The supernatant was discarded before resuspending cells in 500μl of cold FACS media. Acquisition was performed on the BD LSRFortessa cell analyser, and analyzed using FlowJo and GraphPad Prism (**Figure S3**).

For cytotoxic assays the target cells, K562-HLA-A*02:01 were pulsed with the appropriate KRAS associated peptide (10 µg/mL), stained with cell proliferation dye eFluor 450 (eBioscience, 65-0863-14) and then co-cultured with the effector cells (KTCR1 expressing T cells) for 5 hours. Cells were then stained with the fixable viability stain (FVS) - 780 (1/100, 565388, BD Bioscience) and CD8-FITC (1/100, 300905 BioLegend) at 4°C for 30 min. Samples were then washed with 2mL of FACS media and centrifuging for 10 minutes at 400xg at 4°C. The supernatant was discarded and cells resuspended in 500μl of cold FACS media. Acquisition was performed on the BD LSRFortessa cell analyser, and analyzed using FlowJo and GraphPad Prism (**Figure 3b** and **Figure S4**).

### TCR amplification and sequencing

Candidate T cell clones were subjected to alpha-beta TCR amplification and sequencing. We used an adapted version of the template-switch anchored RT-PCR method described previously (*48-49*) where mRNA was extracted from the candidate T cell clones stored in RNAlater (Invitrogen, AM7020) using the RNeasy mini kit (QIAGEN, 74104) following the manufacturer’s protocol. cDNA was generated using a template□switch anchored reverse□transcription polymerase chain reaction (RT□PCR) based on the SMARTer pico cDNA PCR Synthesis Kit (Clontech, Mountain View, CA, USA). The final cDNA product was then subjected to a clean□up step using PCR Clean DX Beads (Aline Biosciences, C-1003). To amplify the TCRα and TCRβ chain, PCR was performed using a pair of primers, that bind to the template switching region (5’□CTAATACGACTCACTATAGGGCAAGCAGTGGTATCAACGCAGAGT and 5’-CTAATACGACTCACTATAGGGC), and the constant region of either the TCRα (5’-AGGCAGACAGACTTGTCACTGGATT) or TCRβ (5’□TCTCTGCTTCTGATGGCTCAAAC) sequence. The PCR assays using NEB enzyme Q5® Hot Start High-Fidelity (NEB, M0491) with the following thermocycling conditions: one cycle at 98°C for 30 sec; 30 cycles at 98°C for 10 sec, 55°C for 10 sec, and 72°C for 20 sec and 1 cycle at 72°C for 5 min. The TCR DNA library from each sample was purified using a 1% agarose e-Gel and a unique band of ∼650 bp in size was excised and extracted using QIAGEN gel extraction kit before being resuspended in RNase/DNase□free water.

A second nested PCR was performed using a nested template switching primer (5’-CGCTCTTCCGATCTCTGGCAGTGGTATCAACGCAGAGTA) and either a TCRα specific primer (5’-TGCTCTTCCGATCTGACCACTGGATTTAGAGTCTCTCAGCTGGT) or TCRβ specific primer (5’□ TCTCTGCTTCTGATGGCTCAAAC). The nested PCR assays were run using the NEB enzyme Q5® Hot Start High-Fidelity (NEB, M0491) as follows: one cycle at 98°C for 30 sec; 10 cycles at 98°C for 10 sec, 65°C for 10 sec, and 72°C for 20 sec and 1 cycle at 72°C for 5 min. The TCR DNA library from each sample was purified using the 1% agarose e-Gel and a unique band of ∼550 bp size was excised and extracted using QIAGEN gel extraction kit before being resuspended in RNase/DNase□free water. The TCRα and TCRβ amplicons were “A-tailed” by adding adenosine to the 3’ end of the amplicons. The amplicons where then ligated pCR4-topo plasmid between the M13F and M13R sequences and transformed into DH5α Competent Cells following the manufactures protocols (Invitrogen, 18265-017). Up to 8 colonies were selected for each TCRα and TCRβ sequence for each of the KRAS^G12V/D^-specific T cell clones and DNA was isolated using a QIAprep Spin Miniprep Kit (QIAGEN, 27104) and sequenced using the M13F and M13R primers. Clone-specific V-gene and J-gene usage, as well as CDR3 sequences were extracted from sequencing data using IMGT/V-QUEST and NCBI-IgBLAST (*42-44*).

### Lentivirus production

To design recombinant TCRs for expression in host T cells, the recovered clonotypes of each KTCR were assembled into a bicistronic alpha-beta gene cassette, which was synthesized de novo and cloned into lentiviral transfer plasmids containing a downstream mStrawberry reporter gene. Replication-incompetent lentiviral particles were then generated to deliver the KTCR gene to healthy donor CD8^+^ T cells. To generate KTCR lentivirus, 80 µg of each transfer plasmid was separately combined with 72 µg of pCMV-ΔR8.91 and 8 µg of pCMV-VSV-G plasmids. The DNA was combined with 430 µL of TransIT-LT1 reagent (Mirus, MIR2305) in a mix made up to 8 mL with OptiMEM (Gibco, 31985062) and incubated for 30 minutes at room temperature. The 8mL DNA/ TransIT-LT1 KTCR mix, was added to eight T75 flasks containing 80-90% confluent 293T/17 [HEK 293T/17] (ATCC® CRL-11268^™^) cells (1mL of transfection mix per flask) along with 9mL of supplemented RPMI-1640 media. Media was then removed 18-20 hours post-transfection and replaced with 5 µl of supplemented RPMI-1640 media. The supplemented RPMI-1640 viral supernatant was then collected at 32, 44, and 56, hours post-transfection. To concentrate virus, supernatants were filtered through a 0.45 µM filter and then centrifuged at 25,000 rpm or 106,750 xg, for 90 minutes at 4°C (Optima XE-90, Beckman-Coulter with SW 32 Ti swinging bucket rotor) to pellet the virus. The virus was then resuspended in 1 mL OptiMEM. Viral titers were determined by testing (in duplicate) 1, 2, 4, 8, 16, or 32 μL of concentrated virus on 1×10^5^ HeLa (ATCC^®^ CCL-2^™^) and K562 cells (ATCC^®^ CCL-243^™^) cells in 24-well format with a final volume of 500 μL of supplemented RPMI-1640 media. Transduction efficiency was determined by measuring the percentage (%) of fluorescent cells detected by flow cytometry. To prepare the cells for flow cytometry analysis, cells were centrifuged for 10 minutes at 400xg at 4°C, the supernatant was discarded and cells were resuspensed in 500μl of cold FACS media with DAPI (1/1000, Sigma D9542). Acquisition was performed on the BD LSRFortessa cell analyser, and analyzed using FlowJo and GraphPad Prism (**Figure S5**).

### Viral transduction

For transduction of CD8^+^ T cells, the appropriate number of PBMCs needed to obtain an MOI of 5 with 100 µL of virus (based on previous titration calculations) were plated in 24-well format and incubated in supplemented RPMI-1640 media, with 300 U/ml rhIL-2 plus 100ng of anti-CD3 and 100ng anti-CD28 soluble antibodies. After 24 hours, 100µL virus was added to the cells. Cells were then incubated for 48 hrs at 37°C, washed, centrifuged at 400 xg for 10 min at room temperature, and cells were resuspended in RPMI-1640 supplemented media with 400 U/ml rhIL-2 and an excess of allogeneic irradiated PBMCs feeders (50 Gy) in a final volume of 0.5 mL. Cells were split every three days by moving 50% of the culture into a new well and adding supplemented RPMI-1640 media with rhIL-2 to give a final rhIL-2 concentration of 400 U/ml in a final volume of 0.5 mL. After 2 weeks of expansion cell aliquots were stored down in cryopreservation media or sorted based on mStrawberry expression via flow cytometry. Sorted cells were expanded again as previously described before being used in the various *in vitro* assays.

The appropriate number of K562 cells needed to obtain an MOI of 5 with 100 µL of HLA-A*02:01-mStrawberry virus (based on previous titration calculations) were plated in 24-well format and incubated in supplemented RPMI-1640 media. After an expansion period, the K562-A*02:01 APCs were sorted based on mStrawberry expression via flow cytometry. Sorted cells were expanded cultured and expanded further before being used in *in vitro* assays.

### Patient derived xenografts

The KTCRs were assessed *in vivo* using a PDX mouse model. Approval was granted by the Animal Care Committee at the University of British Columbia (certificate A19-0035) and the *in vivo* experiments were conducted in a blinded study by the Investigational Drug Program at BC Cancer. A number of KRAS^G12V^ PDX samples had been identified following the methods used by Bournet *et al*. (*51*). DNA was collected from the PDX samples using QIAamp DNA mini kit (Qiagen, 51304) and used in a TaqMan real-time PCR-based allelic discrimination assays (Life Technologies) with custom two allele-specific TaqMan MGB probes which were used to detect KRAS mutational status of the sample (*51*). We analyzed the KRAS^G12V^ positive PDX samples and identified which samples also carried the HLA-A*02:01 allele by following the approach of Browning *et al*. (*52*). CB-17 SCID mice (CB17SC-M, TaconicBiosciences) had the relevant PDX tumour subcutaneously implanted into their lower right back. Once the implanted tumours reached an average size of 100mm^3^, mice received an IV injection of 200μl of PBS containing either 0.5×10^6^ KTCR1 transduced CD8 T cells or 1×10^6^ CD8 T cells, transduced with the mStrawberry-expressing control transfer vector. Tumours were measured using sterilized digital calipers and the volume was calculated using the modified ellipsoid formula Volume (mm^3^) = 1/2(LxW^2^) where length (mm) was the greatest longitudinal measurement taken and the width (mm) the greatest transverse measurement taken. The volume (mm^3^) calculation was converted to a weight (mg) measurement in order to calculate the tumour burden as a percentage of the mouse total weight. We also assessed the effectiveness the KTCR1 treatments by assessing mortality and morbidity during treatments. In particular, we assessed mice for signs of ill health based on body weight loss, change in appetite, behavioural changes such as altered gait, lethargy, tachypnea, pallor, jaundice, and gross manifestations of stress and other signs of ill health.

### Data analysis

Flow cytometry analyses were performed using FlowJo V10.0.8. Sequence data handling was performed using Geneious v8.1.2. Other data handling and statistical analyses were performed using GraphPad Prism software version 8.0.0 for windows (GraphPad Software, California, USA). Results between control groups and test groups were compared using ANOVA analysis and tukey’s multiple comparison test. The Mantel-Cox log rank test was used in the Kaplan-Meier analysis of survival probability analysis of PDX mice. ELISpot’s were analyzed using an AID ELISpot reader and associated software (AID ELISpot Version 7.0).

## End Matter

### Author Contributions and Notes

R.A.H., C.M.R. an S.T. designed the study; G.B.M. and C.S.H. designed the MRM experiments, C.M.R, E.Y., C.S.H., S.D.B., G.S., L.D., N.M.N, C.W., J.H., A.L.M, and T.W., performed the research; J.M.K, D.T.Y., G.B.M. D.J.R.and D.S. and R.A.H provided supervision; R.A.H and C.M.R wrote the manuscript. All authors reviewed and edited the manuscript.

R.A.H., C.M.R., and S.T. are named inventors on patent application PCT/2020/050715 filed by the BC Provinical Health Services Authority.

## Acknowledgments

We thank the BioCanRx network, the BC Cancer Foundation, and the Leon Judah Blackmore Foundation for their ongoing support for this research. We thank the Investigational Drug Program at the BC Cancer Research Institute and the Pancreas Centre BC to the *in vivo* proportion of this body of work. The authors acknowledge the American Association for Cancer Research and its support in the development of the AACR Project GENIE registry as well as members of the consortium for their commitment to data sharing. The MR1 tetramer technology was developed jointly by Dr. James McCluskey, Dr. Jamie Rossjohn, and Dr. David Fairlie, and tetramers for KRAS^G12V^, KRAS^G12D^, and KRAS^WT^ was produced by the NIH Tetramer Core Facility as permitted to be distributed by the University of Melbourne.

## Supplementary

**Table S1:** MRM transition and optimized parameters for the KRAS^WT^ (KLVVVGAGGV) and KRAS^G12D^ (KLVVVGADGV) peptides.

|  | Transitions |  |  |  |  |
| --- | --- | --- | --- | --- | --- |
|  | Product ion | Fragment ion | Dwell time (msec) | Peptide sequence, fragment position, and labeling | Collision energy |
| <u>KRAS<sup>WT</sup></u><br><u>(KLVVVGAGGV)</u> | <u>481.8461</u> | <u>459.2562</u> | <u>45</u> | KRAS-WT.KLVVVGAGGV.+2y6.dimethyl-medium | <u>25</u> |
|  | <u>481.8461</u> | <u>360.1878</u> | <u>45</u> | KRAS-WT.KLVVVGAGGV.+2y5.dimethyl-medium | <u>25</u> |
|  | <u>481.8461</u> | <u>232.1292</u> | <u>45</u> | KRAS-WT.KLVVVGAGGV.+2y3.dimethyl-medium | <u>25</u> |
|  | <u>481.8461</u> | <u>504.4359</u> | <u>45</u> | KRAS-WT.KLVVVGAGGV.+2b4.dimethyl-medium | <u>25</u> |
|  | <u>481.8461</u> | <u>788.5844</u> | <u>45</u> | KRAS-WT.KLVVVGAGGV.+2b8.dimethyl-medium | <u>25</u> |
|  | <u>477.821</u> | <u>459.2562</u> | <u>45</u> | KRAS-WT.KLVVVGAGGV.+2y6.dimethyl-light | <u>25</u> |
|  | <u>477.821</u> | <u>360.1878</u> | <u>45</u> | KRAS-WT.KLVVVGAGGV.+2y5.dimethyl-light | <u>25</u> |
|  | <u>477.821</u> | <u>232.1292</u> | <u>45</u> | KRAS-WT.KLVVVGAGGV.+2y3.dimethyl-light | <u>25</u> |
|  | <u>477.821</u> | <u>496.3857</u> | <u>45</u> | KRAS-WT.KLVVVGAGGV.+2b4.dimethyl-light | <u>25</u> |
|  | <u>477.821</u> | <u>780.5342</u> | <u>45</u> | KRAS-WT.KLVVVGAGGV.+2b8.dimethyl-light | <u>25</u> |
| <u>KRAS<sup>G12D</sup></u><br><u>(KLVVVGADGV)</u> | <u>510.8488</u> | <u>517.2617</u> | <u>45</u> | KRAS-Mut.KLVVVGADGV.+2y6.dimethyl-medium | <u>26.1</u> |
|  | <u>510.8488</u> | <u>418.1932</u> | <u>45</u> | KRAS-Mut.KLVVVGADGV.+2y5.dimethyl-medium | <u>26.1</u> |
|  | <u>510.8488</u> | <u>405.3675</u> | <u>45</u> | KRAS-Mut.KLVVVGADGV.+2b3.dimethyl-medium | <u>26.1</u> |
|  | <u>510.8488</u> | <u>504.4359</u> | <u>45</u> | KRAS-Mut.KLVVVGADGV.+2b4.dimethyl-medium | <u>26.1</u> |
|  | <u>510.8488</u> | <u>603.5044</u> | <u>45</u> | KRAS-Mut.KLVVVGADGV.+2b5.dimethyl-medium | <u>26.1</u> |
|  | <u>510.8488</u> | <u>846.5899</u> | <u>45</u> | KRAS-Mut.KLVVVGADGV.+2b8.dimethyl-medium | <u>26.1</u> |
|  | <u>506.8237</u> | <u>517.2617</u> | <u>45</u> | KRAS-Mut.KLVVVGADGV.+2y6.dimethyl-light | <u>26.1</u> |
|  | <u>506.8237</u> | <u>418.1932</u> | <u>45</u> | KRAS-Mut.KLVVVGADGV.+2y5.dimethyl-light | <u>26.1</u> |
|  | <u>506.8237</u> | <u>397.3173</u> | <u>45</u> | KRAS-Mut.KLVVVGADGV.+2b3.dimethyl-light | <u>26.1</u> |
|  | <u>506.8237</u> | <u>496.3857</u> | <u>45</u> | KRAS-Mut.KLVVVGADGV.+2b4.dimethyl-light | <u>26.1</u> |
|  | <u>506.8237</u> | <u>595.4541</u> | <u>45</u> | KRAS-Mut.KLVVVGADGV.+2b5.dimethyl-light | <u>26.1</u> |
|  | <u>506.8237</u> | <u>652.4756</u> | <u>45</u> | KRAS-Mut.KLVVVGADGV.+2b6.dimethyl-light | <u>26.1</u> |
|  | <u>506.8237</u> | <u>838.5397</u> | <u>45</u> | KRAS-Mut.KLVVVGADGV.+2b8.dimethyl-light | <u>26.1</u> |

**Figure S1:**
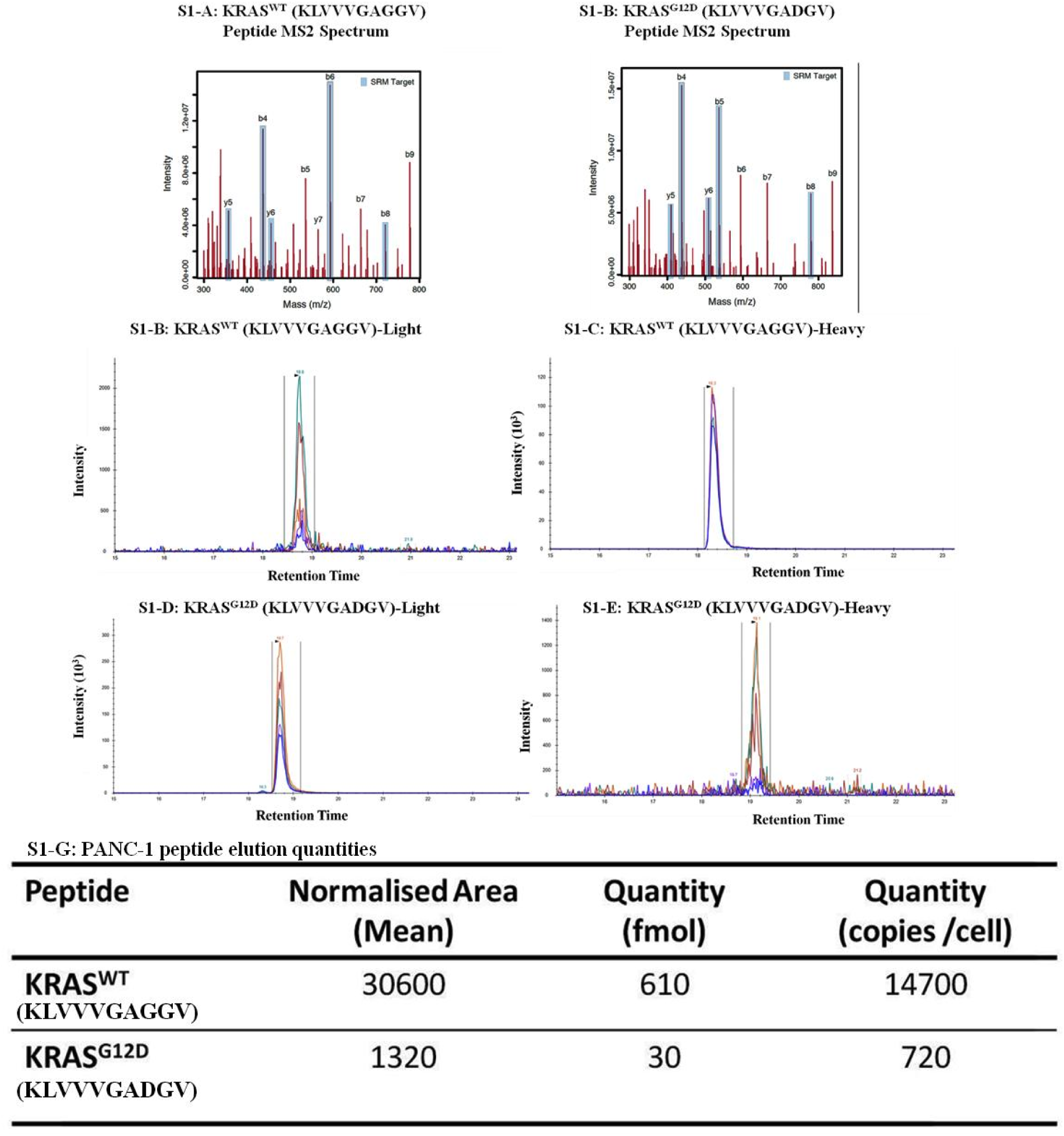
Mass spectrometry analysis of HLA-A*02:01-restricted KRAS^wt^ (KLVVVGAGGV) and KRAS^G12D^ (KLVVVGADGV) aa5-14 peptides eluted from PANC-1 cells. Product-ion peaks for the for KRAS^wt^ and KRAS^G12D^ amino-acid 5-14 peptides, respectively (Figure S1-A, -B). Peak areas for the synthetic peptide standard (Figure S1-C, -D), and the corresponding peak areas for peptide eluted from Panc1 cells (Figure S1-E, -F). The area values and quantities of both KRAS^WT^ and KRAS^G12D^ -specific HLA*02:01-restricted peptides eluted off PANC-1 cells (Figure S1-G). Although it was less abundant than the KRAS^wt^ peptide, the HLA-A*02:01-restricted KRAS^G12D^ peptide can be naturally processed and presented by cells which carry the KRAS^G12D^ mutation and the HLA-A*02:01 allele.

**Figure S2:**
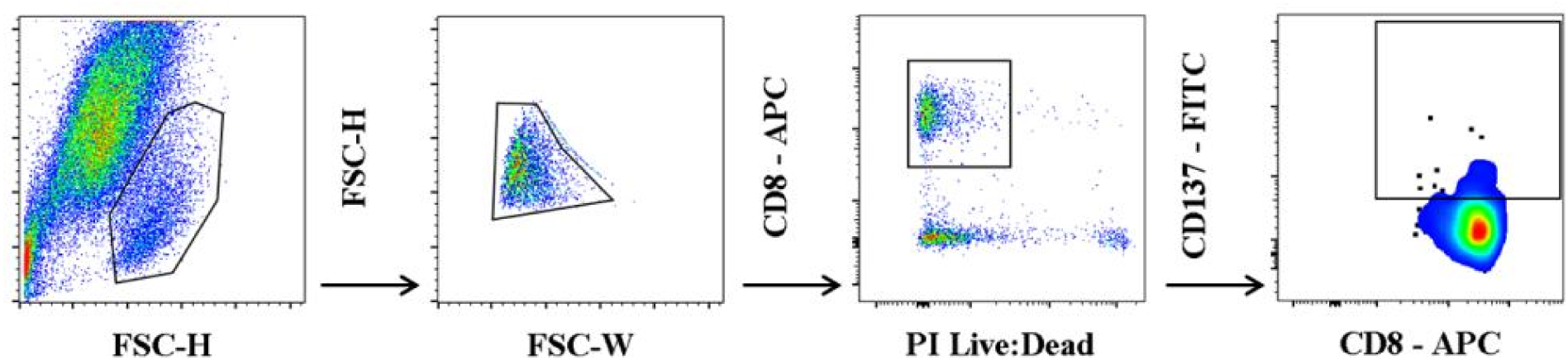
CD137 single cell sort of KRAS^G12D/V/WT^-specific HLA-A*02:01-restricted T cell clones. After debris was eliminated by FSC-A vs. SSC-A gating, cell doublets and clumps were then eliminated by FSC-W vs. FCS-H gating. Live CD8^+^ T cells were determined by gating on the PI negative and CD8^+^ T cells. CD137^+^ expressing CD8^+^ T cells were identified by CD8 vs. CD137 gating and the CD137^+^ CD8^+^ T cells were single cell sorted into 96-well U shaped plate and expanded.

**Figure S3:**
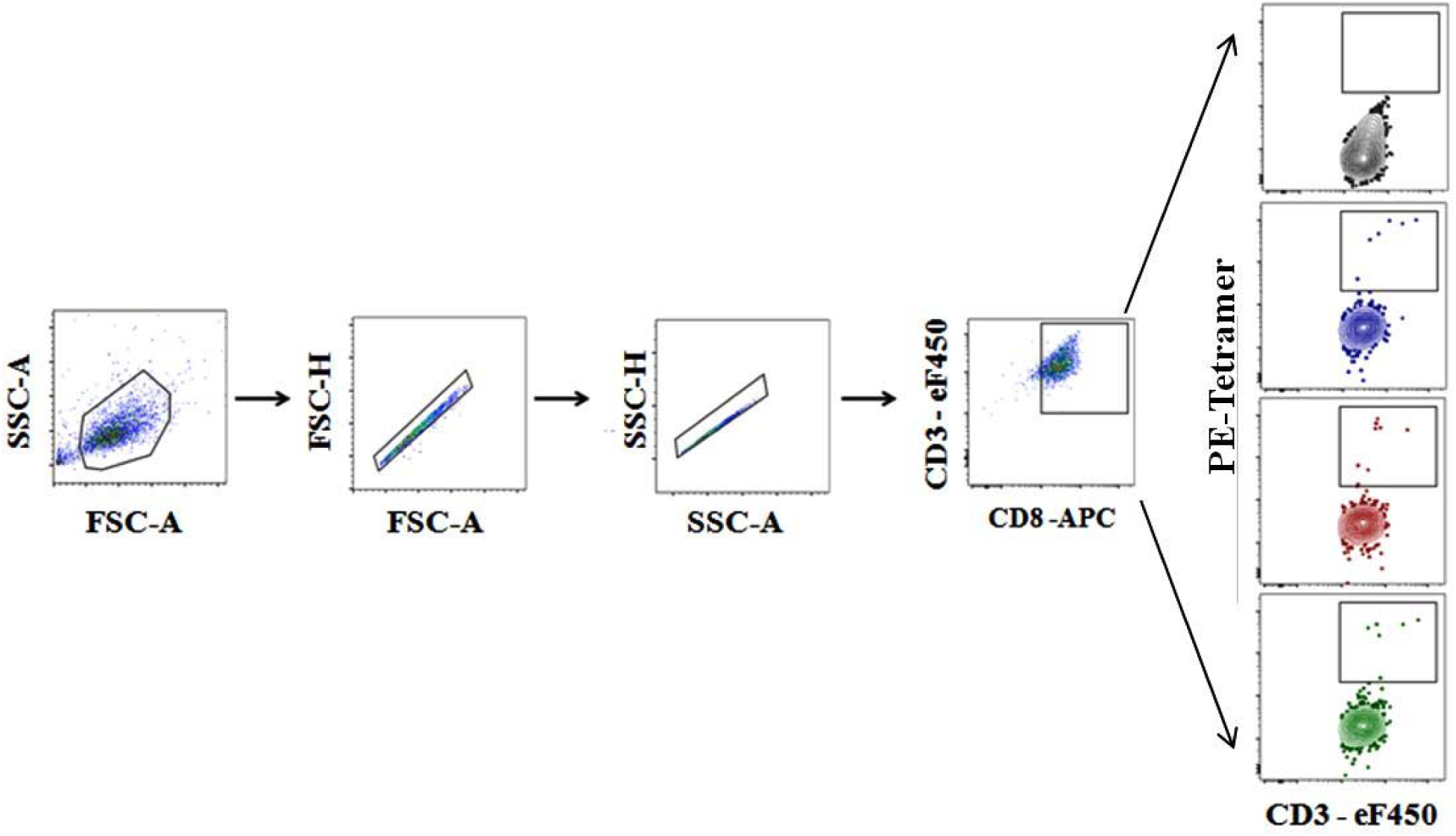
Tetramer staining flow gating protocol. After debris was eliminated by FSC-A vs. SSC-A gating, cell doublets and clumps were then eliminated by FSC-A vs. FSC-H gating followed by SSC-A vs. SSC-H gating. The cells were then gated for CD3 and CD8 expression, followed by CD3 vs. Tetramer gating.

**Figure S4:**
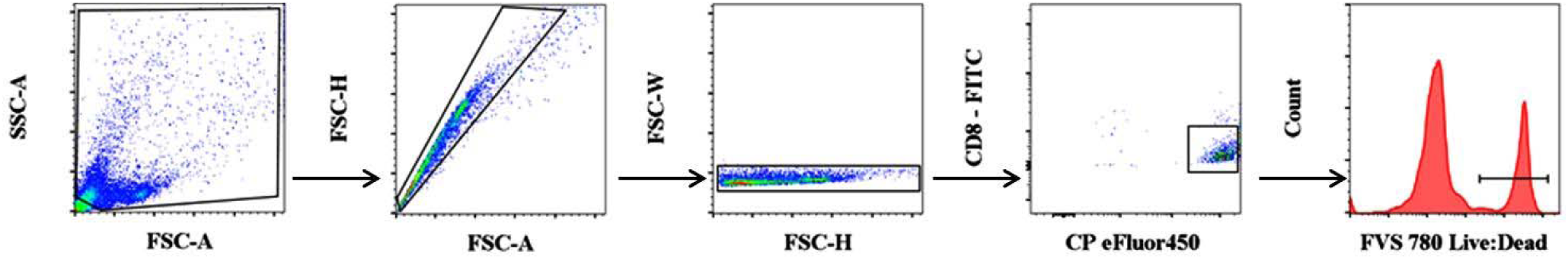
Cytotoxic assay flow gating protocol. After debris was eliminated by FSC-A vs. SSC-A gating, cell doublets and clumps were then eliminated by FSC-A vs. FCS-H gating followed by FCS-H vs. FCS-W gating. The cell proliferation dye eFluor450 stained, peptide pulsed, K562-A*02:01 positive cells were selected for gating them against CD8-FITC stained cells and then determining the percentage of dead cells by gating on FSV780 stained proportion. The flow collected data was analyzed using Flow Jo v10, for windows and GraphPad Prism software version 8.0.0 for windows (GraphPad Software, California, USA).

**Figure S5:**
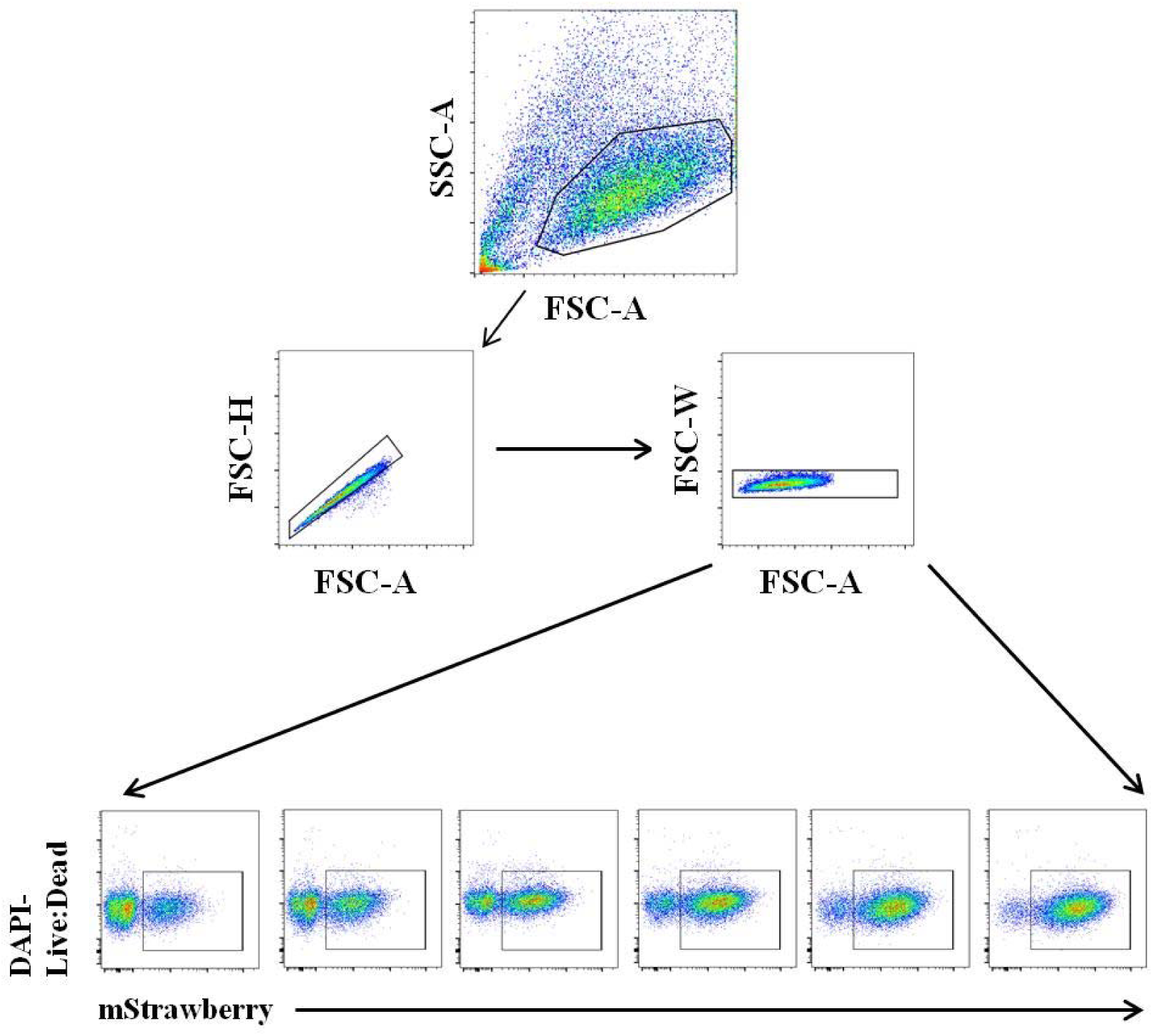
Viral supernatant titration flow gating protocol. After debris was eliminated by FSC-A vs. SSC-A gating, cell doublets and clumps were then eliminated by FSC-A vs. FCS-H gating followed by FCS-A vs. FCS-W gating. The live cells were determined by gating on the DAPI negative cells and on mStrawberry to assess the level of mStrawberry expression within the cells in order to measure transduction efficiency. The flow collected data was analyzed using Flow Jo v10, for windows and GraphPad Prism software version 8.0.0 for windows (GraphPad Software, California, USA).

